# PTEN and BRCA1 tumor suppressor loss associated tumor immune microenvironment exhibits differential response to therapeutic STING pathway activation in a murine model of ovarian cancer

**DOI:** 10.1101/2022.06.27.497846

**Authors:** Noor Shakfa, Deyang Li, Gwenaëlle Conseil, Elizabeth Lightbody, Juliette Wilson-Sanchez, Ali Hamade, Stephen Chenard, Brian Laight, Afrakoma Afriyie-Asante, Kathrin Tyryshkin, Martin Köbel, Madhuri Koti

**Author notes:** **CORRESPONDING AUTHOR:** Madhuri Koti, *DVM, MVSc, PhD*, Department of Biomedical and Molecular Sciences & Obstetrics and Gynecology, Queen’s University, Kingston, Ontario, K7L3N6, Canada, *e-mail:.

## Abstract

**Background:** High grade serous ovarian carcinoma (HGSC) is the most lethal gynecologic malignancy characterized by chemoresistance and high rates of recurrence. HGSC tumors display a high prevalence of tumor suppressor gene loss. Loss of *BRCA1* and *PTEN* function due to mutations or epigenetic influence have been widely associated with variable clinical outcomes, where tumors with *BRCA1* mutations exhibit increased chemosensitivity and those with *PTEN* mutations have been reported to exhibit chemoresistance. Given the established type 1 interferon regulatory function of *BRCA1* and *PTEN* genes and associated contrasting T cell infiltrated and non-infiltrated tumor immune microenvironment (TIME) states, in this study we investigated the potential of Stimulator of Interferon Genes (STING) pathway activation in improving overall survival via enhancing chemotherapy response, specifically in tumors with *PTEN* deficiency.

**Methods:** Expression of PTEN protein was evaluated in tissue microarrays generated using pre-treatment tumors collected from a cohort of 110 patients with HGSC. Multiplex immunofluorescence staining was performed to determine spatial profiles and density of selected lymphoid and myeloid cells. *In vivo* studies using the syngeneic murine HGSC cell lines, ID8-*Trp53*^*-/-*^; *Pten*^*-/-*^and ID8-*Trp53*^*-/-*^; *Brca1*^*-/-*^, were conducted to characterize the TIME and response to carboplatin chemotherapy in combination with exogenous STING activation therapy.

**Results:** Tumors with absence of PTEN protein exhibited a significantly decreased disease specific survival and intra-epithelial CD68+ macrophage infiltration as compared to intact PTEN expression. *In vivo* studies demonstrated that *Pten* deficient ovarian cancer cells establish an immunosuppressed TIME characterized by increased proportions of M2-like macrophages, GR1+ MDSCs in the ascites, and reduced effector CD8+ cytotoxic T cell function compared to *Brca1* deficient cells; further, tumors from mice injected with *Pten* deficient ID8 cells exhibited an aggressive behavior due to suppressive macrophage dominance in the malignant ascites. In combination with chemotherapy, exogenous STING activation resulted in longer overall survival in mice injected with *Pten* deficient ID8 cells, reprogrammed intraperitoneal M2-like macrophages derived from *Pten* deficient ascites to a M1-like phenotype and rescued CD8+ cytotoxic T cell activation.

**Conclusions:** This study reveals the importance of considering the influence of cancer cell intrinsic genetic alterations on the TIME for therapeutic selection. We establish the rationale for the optimal incorporation of interferon activating therapies as a novel combination strategy in PTEN deficient HGSC.

## INTRODUCTION

High-grade serous ovarian carcinoma (HGSC) is the most common and aggressive histological subtype of epithelial ovarian cancer. Patients with HGSC often present with advanced disease, generally managed with debulking surgery followed by platinum and taxane-based combination chemotherapy^1^. Although most patients show initial sensitivity to chemotherapy, over 80% relapse and subsequently develop resistance to platinum^1^. Despite the success of poly-ADP ribose polymerase (PARP) inhibitors in a subpopulation of HGSC patients (homologous recombination deficient and BRCA-mutated), newer treatment options such as anti-angiogenic agents and immune checkpoint inhibitors have demonstrated modest or no improvements in HGSC^2,3^. While significant advancements have been made to delineate pre-treatment tumor immune states that associate with differential treatment outcomes, their therapeutic vulnerabilities remain to be fully exploited. Our previous reports on the pre-treatment tumor immune microenvironment (TIME) states in patients with HGSC have demonstrated that higher expression of type I interferon genes (IFN-1) associates with increased response to chemotherapy ^4,5^.

Recent evidence further suggests that the evolution of distinct TIME states and downstream responses can be driven by cancer cell intrinsic alterations in tumor suppressor genes that regulate cellular IFN-1 pathways^6^. While mutations in the tumor suppressor gene, *TP53*, are universal and a characteristic feature of HGSC, additional mutations in genes associated with DNA damage repair (DDR) pathways are present in approximately 50% of HGSC cases^7^. Among the DDR genes, mutations in breast cancer type I susceptibility protein 1 and 2 (*BRCA1/BRCA2*) have been widely reported to correlate with increased chemosensitivity^8,9^. Interestingly, tumors with DDR deficiency also exhibit high infiltration of CD8+ T cells^10^. In contrast to the chemosensitive behavior of tumors with *BRCA1* mutations, those with mutations in the phosphatase and tensin homolog (*PTEN*) gene (present in ∼10% of HGSC cases) frequently exhibit a chemoresistant profile^7^. In other solid tumors, *PTEN* deficiency via mutations, copy number loss, or epigenetic silencing is also associated with a decreased infiltration of immune cells in the tumor microenvironment^11^. This can potentially be attributed to *PTEN* deficiency associated cytokine signaling that stimulates an immunosuppressive microenvironment^11^. Importantly, both *BRCA* and *PTEN* genes have a regulatory function in activation of IFN-1 pathways^12,13^. While therapies such as PARP inhibitors have shown promise in *BRCA1/2* and other homologous recombination (HR) deficient immune infiltrated HGSC tumors^14,15^, there remains an unmet need for those patients who exhibit an underactive immune state that often accompanies a chemoresistant tumor phenotype. In the context of chemoresistance associated genetic alterations, reports in melanoma and glioma have shown that *PTEN* loss associated PI3K/AKT pathway activation plays a crucial role via altering macrophage activation within the TIME towards a tumor promoting functional state^16,17^.

We have previously shown that therapeutic activation of STING pathway leads to production of chemokines such as CXCL10 and CCL5 which recruit cytotoxic CD8+ T cells to the TIME, enhances response to carboplatin chemotherapy, sensitizes to PD-1 immune checkpoint blockade and overall survival in the ID8 murine model of HGSC^18^. Here, we aimed to investigate whether cancer cell intrinsic loss of PTEN or BRCA1 influence the TIME, response to chemotherapy, and survival outcomes. We further provide evidence supporting the benefit of exogenously activating the STING pathway in the context of BRCA1 or PTEN deficiency. Findings from this study demonstrate that although therapeutic STING pathway activation leads to improved response to chemotherapy and improved overall survival, PTEN deficient tumors with a suppressed pre-treatment TIME state exhibit a more pronounced benefit through re-invigoration of the TIME.

## METHODS

### Patient Specimens

Archival tumor specimens from 110 HGSC patients were accessed under the institutional health research ethics board approval (University of Calgary, AB, Canada). Tissue microarrays (TMAs) were constructed from formalin-fixed paraffin-embedded tumors following histopathological confirmation of HGSC by a pathologist (Dr. M. Köbel). A total of 110 chemo naïve tumors were used for subsequent analysis. The clinicopathological parameters of the cohort are provided in Supplemental Table 1. Duplicate or triplicate 0.6 mm cores were extracted from the areas of interest of each tumour specimen and embedded into a recipient block of paraffin for TMA construction.

### Immunostaining of HGSC TMA

Four μm TMA sections were stained with PTEN Rabbit mAb (1/50 dilution; clone D4.3 XP; Cell Signaling Technologies, MA, USA). Multiplex immunofluorescence staining of TMA sections was performed using antibodies specific to CD8, CD68, and pan-cytokeratin (panCK) (Molecular and Cellular Immunology Core, BC Cancer Research, BC, Canada). Slides were scanned using Vectra Polaris (MOTiF™) Multispectral Imaging System (Akoya Biosciences, MA, USA). Individual tumor cores on the scanned TMA cores were segmented and annotated for stromal vs. epithelial compartments using the Halo® Link image analysis Software (Indica Labs, NM, USA). Algorithms to detect markers of interest were generated on Halo® Link outputting both a total cell number using DAPI staining and total cells positively stained with markers of interest within tissue compartments for PTEN deficient and PTEN intact tumors. Patient clinical data from samples within TMAs were accessed for association with clinical parameters.

### Murine ovarian cancer cell lines

The syngeneic murine ovarian cancer cell lines ID8-*Trp53*^*-/-*^; *Pten*^*-/-*^and ID8-*Trp53*^*-/-*^; *Brca1*^*-/-*^, were kindly provided by Dr. Ian McNeish (Imperial College, UK; Supplemental Figure 2C). Both cell lines were maintained in Dulbecco’s Modified Eagle’s Medium (Sigma-Aldrich, ON, Canada) supplemented with 4% heat-inactivated fetal bovine serum (Sigma-Aldrich, ON, Canada), 1% penicillin-streptomycin (100 μg/mL), and 1% insulin transferrin sodium selenite (ITS) liquid media supplement (Sigma-Aldrich, ON, Canada) and incubated at 37°C/5% CO_2_. Cells for bioluminescent imaging were fluorescently tagged with firefly luciferase by viral transduction using previously established protocols^19^.

### IC50 of carboplatin using clonogenic assay

The IC50 of carboplatin was determined using clonogenic assays as per previously established methods^20^. ID8 *Trp53*^*-/-*^; *Pten*^*-/-*^and ID8 *Trp53*^*-/-*^; *Brca1*^*-/-*^cells were seeded at density of 750 cells/well in a 6-well plate and allowed to adhere for 8 hours. Supernatants were replaced with fresh complete DMEM containing serial dilutions (0-250 μM) of carboplatin (10 μg/mL) or media control.

Cells were incubated at 37°C/5% CO_2_ for ∼4 days, or until colonies containing approximately 50 cells formed. Cells were fixed using 4% paraformaldehyde (PFA) diluted in 1X phosphate buffered saline (PBS) and stained with 0.25% crystal violet in 80% methanol. Colonies were imaged and quantified to generate a Percent Area (PA) value using ImageJ Software®. Dose-response curves and IC50 concentrations were computed on GraphPad Prism v9.0 software (GraphPad Software, CA, USA)

### Macrophage migration assay

IC21 murine peritoneal macrophage cell line was obtained from American Type Culture Collection (ATCC; Manassas, VA, USA) and cultured in RPMI 1640 (Sigma-Aldrich, ON, Canada) supplemented with 10% FBS and 1% penicillin/streptomycin. ID8 *Trp53*^*-/-*^; *Pten*^*-/-*^and ID8 *Trp53*^*-/-*^; *Brca1*^*-/-*^cells were resuspended at a density of 5×10^5^ in 4 mL complete growth media and seeded in 6 well plates. Conditioned media from both lines were collected 72 h post seeding. Migration assays were performed in uncoated 24-well transwell plates with 8 μm pore inserts and 6.5 mm in diameter (Corning, NY, USA). IC21 cells were added at a cell concentration of 5 × 10^5^ in 100 μL of serum-free media into the upper chamber and allowed to migrate through the insert membrane for 16 hours in a 37°C/5% CO_2_ atmosphere. Conditioned media (600 μL) from either ID8 derivative cell line was placed into the lower chambers. Recombinant murine macrophage chemoattractant protein 1 (MCP-1) (20 ng/mL; Peprotech, NJ, USA) was used as a positive control. Cells which migrated onto the transwell inserts were fixed with 4% PFA, stained with DAPI (1 μg/ml; Sigma-Aldrich, ON, Canada) and mounted onto a positively charged slide. Inserts were imaged using Evos® Cell Imaging System and quantified using ImageJ software®.

### Immunofluorescence staining to detect cytosolic DNA in ID8 cells

Immunofluorescence staining was performed to assess the constitutive cytosolic dsDNA expression in untreated ID8 cells. ID8-*Trp53*^*-/-*^; *Pten*^*-/-*^or ID8-*Trp53*^*-/-*^; *Brca1*^*-/-*^cells were seeded at a density of 5 × 10^5^ on coverslips placed in a 6-well plate and left to adhere overnight in complete growth media at 37°C/5% CO_2_. Coverslips were washed with PBS and fixed for 10 minutes with 4% PFA. Following permeabilization of cells (0.2% TritonX-100 in 1X PBS) and blocking (0.1% TritonX-100, 1% BSA in 1X PBS) at room temperature, cells were incubated in anti-dsDNA monoclonal antibody (1:100; EMD Millipore, ON, Canada) overnight at 4°C. Coverslips were washed with blocking solution 3 times at room temperature and incubated in blocking buffer containing Alexa Flour 488 anti-mouse IgG antibody (1:300; Invitrogen, MA, USA) for 1 h at room temperature in the dark. After counterstaining with DAPI for 10 min, coverslips were rinsed with PBS and mounted with antifading fluorescence medium (Invitrogen, Massachusetts, USA) onto a slide. Slides were imaged using Evos® Cell Imaging System and quantified using ImageJ software®.

### Western Blotting

Immunoblotting analyses were performed by briefly washing ID8-*Trp53*^*-/-*^; *Pten*^*-/-*^and ID8-*Trp53*^*-/-*^; *Brca1*^*-/-*^cells three times with cold PBS and lysing in RIPA buffer containing protease and phosphatase inhibitors (Thermofisher Scientific, ON, Canada). The protein concentrations of cell lysates were quantified using a Bradford assay with bovine serum albumin (BSA) standards. Proteins were denatured by heating samples for 10 mins at 95°C. Equal amount of total protein (25 μg/well) was resolved by electrophoresis on 8% Tris-Acrylamide gels and transferred onto 0.2 μm nitrocellulose membranes (Bio-Rad Laboratories, CA, US). Membranes were blocked for 1 h at room temperature in 5% skim milk diluted in Tris-buffered saline supplemented with 0.1% Tween 20 (TBST). Membranes were incubated in primary antibodies for PTEN (Cell Signaling Technology, MA, USA), BRCA1 (Sigma Aldrich, ON, Canada), and STING (Cell Signaling Technology, MA, USA) at a dilution of 1:1000 in 5% skim milk and 1x TBST at 4°C overnight. Nitrocellulose-bound primary antibodies were detected with anti-rabbit IgG horseradish peroxidase-linked antibody using the Clarity Western Peroxide and Luminol/Enhancer reagents (Bio-Rad Laboratories, CA, USA).

### In vivo studies

All *in vivo* procedures performed were approved by the University Animal Care Committee at Queen’s University. A total of 5-6 × 10^6^ ID8-*Trp53*^*-/-*^; *Pten*^*-/-*^or ID8-*Trp53*^*-/-*^; *Brca1*^*-/-*^cells in PBS were injected via intraperitoneal route in 8–10-week-old female C57BL/6 mice (Charles River Laboratories International Inc.). All mice were maintained under specific pathogen-free conditions. Treatments were initiated three weeks post-cancer cell injections. Mice were randomised into three treatment groups (n = 10-15 per group): vehicle (PBS), carboplatin, or carboplatin + STING agonist (2’3’=c-di-AM(PS)2 (Rp, Rp); InvivoGen, CA, USA). Carboplatin was used at a dose of 10 mg/kg twice/week for 4 consecutive weeks and STING agonist was used at a dose of 4 mg/kg once/week for 3 doses (Figure 5A). To determine the effect of cancer cell intrinsic vs immune cell intrinsic STING activation *in vivo*, 5-6 × 10^6^ ID8-*Trp53*^*-/-*^; *Pten*^*-/-*^or ID8-*Trp53*^*-/-*^; *Brca1*^*-/-*^cells in PBS were injected via intraperitoneal route in 8–10-week-old female C57BL/6J-*Sting1*^*gt*^/J (STING-KO) mice (Jackson Laboratories International Inc., CT, USA) and profiled at endpoint.

### *In vivo* bioluminescence imaging

A total of 5-6 × 10^6^ ID8-*Trp53*^*-/-*^; *Pten*^*-/-*^or ID8-*Trp53*^*-/-*^; *Brca1*^*-/-*^luciferase-tagged cells in PBS were injected via intraperitoneal route in 8–10-week-old female C57BL/6 mice (Charles River Laboratories International Inc.). Imaging was started at day-7 post-cell injection, performed once a week, until week 4 in order to record the signal emitted over tumor and metastasis progression. Intraperitoneal injection of D-Luciferin firefly (Perkin Elmer, France) at a dose of 150 mg/kg body weight was performed on animals. Imaging and data processing were performed using the IVIS Spectrum imager equipped with the Living Image® 4.3.2 software (Perkin Elmer, France). Acquisition began 10 min after injection with D-Luciferin, with mice under gas anesthesia (2% isoflurane). Images were measured between 500 and 660 nm emission wavelengths.

### Multiplex cytokine analysis of post-treatment plasma and ascites fluid

Ascites fluid and plasma samples collected 24 h following initial STING agonist dose or post carboplatin chemotherapy alone from each treatment group were subjected to multiplex cytokine analysis using the mouse Cytokine Array/Chemokine Array 31-Plex Discovery Assay (includes Eotaxin, G-CSF, GM-CSF, IFN gamma, IL-1alpha, IL-1beta, IL-2, IL-3, IL-4, IL-5, IL-6, IL-7, IL-9, IL-10, IL-12 (p40), IL-12 (p70), IL-13, IL-15, IL-17A, IP-10 (CXCL10), CXCL1, LIF, LIX, MCP-1 (CCL2), M-CSF, MIG (CXCL9), MIP-1alpha, MIP-1beta, MIP-2, RANTES (CCL5), TNF alpha, and VEGF) at Eve Technologies (AB, Canada). All samples were analysed in biological triplicates. The standard curve regression was used to calculate the concentration of each target cytokine.

### Local and systemic immune profiling using polychromatic flow cytometry

Measurement of immune cell proportions within splenocytes and ascites cells was conducted using flow cytometry to characterize systemic and local immune profiles, respectively. Splenocytes from all 3 treatment groups (control, carboplatin, carboplatin + STING agonist) were collected 24 h post initial carboplatin or carboplatin + STING agonist dose. Following sacrifice of mice, ascites was aspirated using an 18-gauge needle. Single-cell suspensions of splenocytes were prepared by mechanical dissociation of the tissue followed by passing dissociated tissue through a 40 μm cell strainer. 1x red blood cell (RBC) lysis buffer was added to ascites cells and splenocytes. Cells were digested in RPMI-1640 media containing 20 μg/mL DNase (Roche, Basel, Switzerland) and 1 mg/mL collagenase IV (StemCell Technologies, BC, Canada) for 30 min at 37°C/5% CO_2_. Cells were washed in PBS, counted and subjected to flow cytometry analysis of target immune populations (CD45+ cells) using a lymphoid panel (CD3 T cells, CD4 helper T cells, CD8 cytotoxic T cells, CD19 B cells, NK1.1 natural killer cells, CD62L/CD69 T cell activation, PD-1 checkpoint; Biolegend, CA, USA) and myeloid panel (CD11b myeloid cells, CD11c dendritic cells, F4/80 macrophages, CD80 M1-like macrophages, CD206 M2-like macrophages, GR1 myeloid derived suppressor cells, PD-L1 checkpoint), on the CytoFlex-S Flow Cytometer (Beckman Coulter, ON, Canada). Single colour positive controls, as well as unstained and fluorescence-minus-one (FMO) negative controls were used for each antibody and their respective panel to determine gates. Gating and analysis were conducted using FlowJo® v10 Software (BD, ON, Canada).

### NanoString based tumour immune transcriptomic profiling

Immune gene expression profiling was performed using the pre-built nCounter Mouse PanCancer Immune Profiling panel which includes genes associated with immune function, various cancer-related pathways, and housekeeping genes (NanoString Technologies, WA, USA) to determine both the baseline differences in the TIME generated from ID8 cells of different genotypes and to measure the effect of STING agonist treatment on the TIME. Total RNA of tumors collected from mice injected with ID8-*Trp53*^*-/-*^; *Pten*^*-/-*^or ID8-*Trp53*^*-/-*^; *Brca1*^*-/-*^cells with and without treatment was isolated at endpoint using the total RNA Purification Kit (Norgen Biotek Corporation, ON, Canada) as per the manufacturer’s instructions. The purity and concentration of isolated RNA was estimated spectrophotometrically using the NanoDrop ND-100 spectrophotometer (NanoDrop Technologies, DE, USA). Total RNA (150 ng) was used as template for digital multiplexed profiling at the Queen’s Molecular Pathology Laboratory’s (QLMP) as per previously established protocols^18^. Raw nCounter NanoString counts were normalized using nSolver software 3.0 (NanoString Technologies, WA, USA), using NanoString’s built-in positive controls. mRNA content normalization was performed using housekeeping genes and overall assay efficiency was calculated using the geometric mean of each control. Differential gene expression between comparison groups was computed on nSolver, and Benjamini-Hochberg method was used to adjust for the false discovery rate. Heatmaps were generated on nSolver software (NanoString, WA, USA.

### Macrophage Polarization Assay

IC21 cells were seeded at a density of 5×10^5^/well in a 6-well plate and incubated overnight at 37°C in complete media. Ascites generated from untreated mice injected with either ID8-*Trp53*^-/-^; *Pten*^-/-^or ID8-*Trp53*^-/-^; *Brca1*^-/-^cells was centrifuged to remove cellular debris and fluid fraction was collected separately. Ascites fluid was passed through Amicon® centrifugal filter concentrators (EMD Millipore, ON, Canada). Concentrated ascites fluid protein levels were quantified using a Bradford assay. IC21 cells were incubated in media containing equal concentrations of ascites fluid in media from either genotype (1:100 ratio) for 24 h. Expression of M2-like macrophage associated markers (F4/80, CD206 and PDL1) was measured using flow cytometry.

### Macrophage and T-cell co-culture assay

Splenocytes were collected from a healthy female mouse spleen, mechanically dissociated, passed through a 40 μm strainer and followed by RBC lysis. Single cell suspensions were then incubated in RPMI-1640 supplemented with 20 μg/mL DNase and 1 mg/mL collagenase IV for 30 min at 37°C/5% CO_2_. Splenic T cells were enriched using a magnetic-based commercial CD8 T cell negative selection kit (StemCell Technologies, BC, Canada; Supplemental Figure 3A) as per the manufacturer’s protocol. Ascites from mice (n=5 for each group) injected with either ID8-*Trp53*^*-/-*^; *Pten*^*-/-*^or ID8-*Trp53*^*-/-*^; *Brca1*^*-/-*^cells were collected, passed through a 70 μm strainer, lysed using 1x RBC lysis buffer and incubated in RPMI-1640 supplemented with 20 μg/mL DNase and 1 mg/mL collagenase IV for 30 min at 37°C/5% CO_2_. Macrophages were enriched using a magnetic-based commercial F4/80 macrophage positive selection kit (Miltenyi Biotec, MD, USA; Supplemental 3B) as per manufacturer’s instructions. A 96-well plate was coated with anti-CD3e (1 mg/mL; Thermofisher, CA, USA) and anti-CD28 (5 mg/mL; Thermofisher) antibodies for 4 h at 37°C/5% CO_2_. CD8+ T cell-to-macrophage ratio was seeded at 1:1 (100,000 of each cell type) into triplicate wells for each mouse in 200 μL of IL-2 (5 IU) treated media containing 1% gentamycin and 10% FBS and incubated for 48 h at 37°C/5% CO_2_. Four hours prior to the 48-hour timepoint, brefeldin A (Cayman, Michigan, USA) was added to each well (10 μg/mL). Supernatant containing T cells in suspension was aspirated and placed into a new 96-well plate for flow cytometric analysis of activation state stained with anti-CD45, CD3, CD8, F480, PD-1 and IFNγ antibodies (Biolegend). 1x EDTA was used to lift adherent macrophages off of wells and cells were placed into a new 96-well plate for staining of macrophage state. The expression of CD45, F480, CD80, CD206, CD8 and PD-L1 (Biolegend, CA, USA) was measured using polychromatic flow cytometry (Beckman Coulter). Intracellular levels of IFNγ in CD8+ T cells treated with macrophages from different genotypes was measured to determine the influence of macrophages on T cell activation.

### Statistical Analysis

Statistical analyses were performed using GraphPad Prism v9.0 software (GraphPad Software) as described in the results. Statistics for Kaplan Meier survival analysis on patient cohort was performed using the survminer survival package in R (version 3.5.2, R Studio, MA, USA). All analysis used Mann-Whitney nonparametric test (for data that deviates from normality) to compare two conditions unless otherwise indicated. Results are expressed as a mean ±SD. A p-value <0.05 was considered statistically significant.

## RESULTS

### PTEN deficient human HGSC tumors associate with decreased overall survival and exhibit a distinct macrophage infiltration profile

Pre-chemotherapy treated tumors from 110 patients with HGSC (Supplemental Table 1) were evaluated for PTEN protein expression using a pre-established PTEN scoring system^21^. Loss of PTEN protein expression was observed in 9.1% of cases within this cohort. Tumors with either complete absence of PTEN protein expression (n=10) in epithelial compartments or with normal expression (n=57) were subjected to further analyses. Kaplan-Meier survival analysis demonstrated a statistically significant shorter disease specific survival (DSS; log-rank test, p = 0.0376; HR = 0.45; Figure 1A) in tumors with absence of PTEN expression compared to those with normal PTEN expression. Immunofluorescence staining revealed decreased CD8+ cytotoxic T cells and CD68+ macrophages in tumors with complete absence of PTEN expression (Figure 1B) compared to tumors with PTEN presence (Figure 1C), however, these differences were not statistically significant potentially due to Intratumoral heterogeneity. In this cohort, CD68+ macrophages were only observed to heavily infiltrate the stromal compartment of PTEN deficient tumors and displayed low densities in the epithelial compartments of these tumors (Figure 1D; p = 0.0005 between compartments). In contrast, PTEN intact tumors displayed intra-epithelial CD68+ macrophages revealing macrophage patterns specific to the absence of PTEN (p>0.05 between compartments). This suggests a potential relationship between HGSC PTEN alterations driving variations in intratumoral macrophage infiltration patterns.

**Figure 1.**
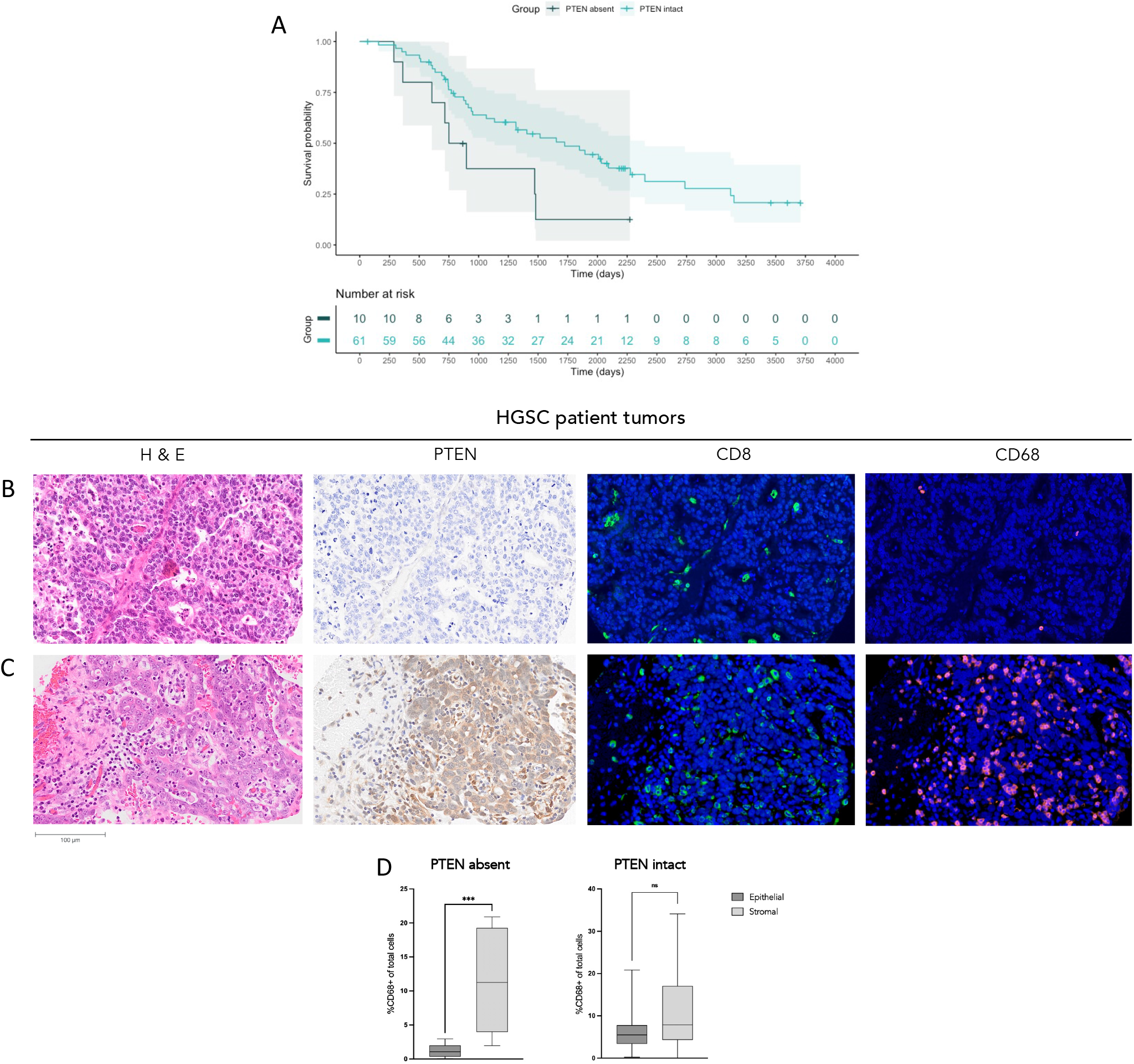
PTEN deficiency in the tumor immune microenvironment of HGSC patient samples leads to shorter overall survival and decreased immune infiltration. (A) Kaplan-Meier survival analysis using clinical data for patients with PTEN intact (n=10) had significantly higher disease specific survival (DSS) than patients with PTEN absent (n=57; p=0.0376). Fluorescent immunostaining of CD8+ cytotoxic T cells and CD68+ macrophages within tumors of patients with (B) complete absence of PTEN (PTEN absent; n=9) and (C) complete presence of PTEN (PTEN intact; n=51) in stromal and epithelial compartments of patient tumors using immunohistochemistry. Immunofluorescent staining of CD8 and CD68 expressing cells were quantified in different tissue compartments. (D) Infiltration patterns of CD68+ macrophages between different compartments in tumors with PTEN absence and PTEN intact were compared, revealing a significantly higher amount of CD68+ macrophages in the stroma compared to the epithelium in tumors with PTEN absence (p=0.0005). Average of duplicate or triplicate cores for each sample was taken and Mann-Whitney non-parametric test was used to determine statistical significance of immune cell infiltration. Log-rank test was applied to determine statistical significance of Kaplan-Meier survival analysis, using R statistical software. ** p<0.005, *** p<0.001, **** p<0.0001. P-value <0.05 was considered statistically significant. ns: not significant.

### Cancer cell genotype influences the TIME and disease progression in the ID8 murine model of HGSC

We first evaluated whether the cancer cell intrinsic loss of *Pten* or *Brca1* led to survival differences in the ID8 syngeneic murine model of HGSC. Mice were injected with ID8-*Trp53*^*-/-*^, ID8-*Trp53*^*-/-*^; *Pten*^*-/-*^, *or* ID8-*Trp53*^*-/-*^; *Brca1*^*-/-*^cells (henceforth denoted as ‘*Trp53* deficient’; ‘*Pten* deficient’ or ‘*Brca1* deficient’ cells, respectively). Mice injected with *Pten* deficient cells had a statistically significant shorter median OS of 39 days compared to those injected with *Brca1* deficient cells, which displayed a median OS of 45 days (*p*=0.0002), both genotypes displaying a shorter OS than mice injected with *Trp53* deficient ID8 cells (56 days; Figure 2A).

**Figure 2.**
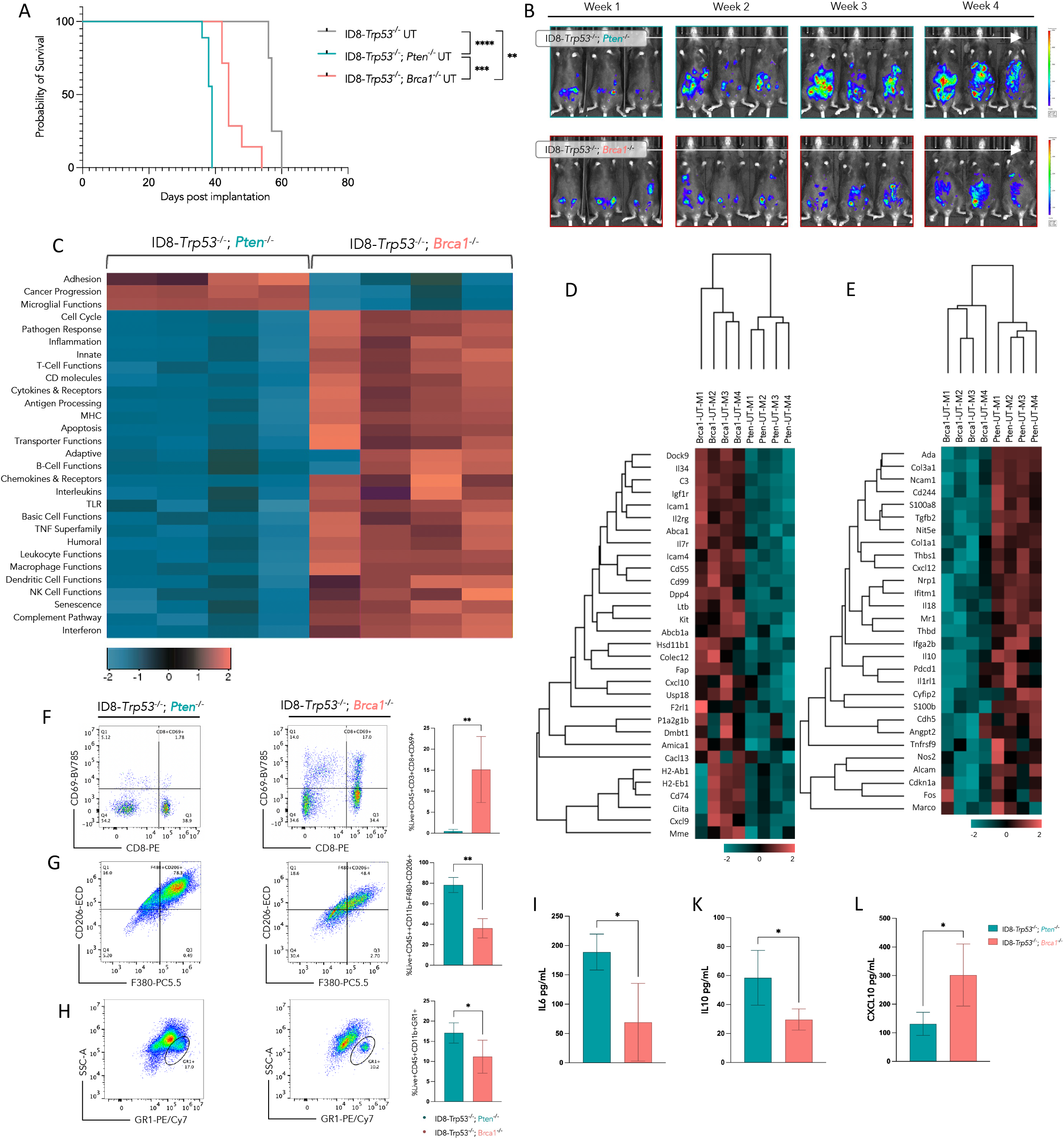
Pten deficient HGSC TIME and progression is distinct from Brca1 deficient HGSC. (A) Kaplan-Meier survival analysis of mice injected with either ID8-*Trp53*^-/-^, ID8-*Trp53*^-/-^; *Pten*^-/-^and ID8-*Trp53*^-/-^; *Brca1*^-/-^cells. (n = 10-15) for each treatment group. (B) *In vivo* bioluminescent imaging of mice injected with ID8-*Trp53*^-/-^; *Pten*^-/-^and ID8-*Trp53*^-/-^; *Brca1*^-/-^luciferase-tagged cells tracking tumor progression once/week over 4 weeks. Total RNA from untreated tumors of different genotypes was subject to NanoString gene expression profiling using the panCancer immune gene panel displayed as a (C) heat map showing differential expression pattern of groups of genes involved in various immune functions, (D) 2-fold differentially expressed genes and (E) 0.5-fold differentially expressed genes. Proportion of cells derived from ascites of mice injected with ID8-*Trp53*^-/-^; *Pten*^-/-^and ID8-*Trp53*^-/-^; *Brca1*^-/-^cells expressing (F) CD45+CD3+CD8+CD69+, (G) CD45+CD11b+F4/80+CD206+, and (H) CD45+CD11b+GR1+. (I) IL-6, (K) IL-10 and (G) CXCL-10 cytokine levels derived from the ascites fluid of untreated mice injected with ID8-*Trp53*^-/-^; *Pten*^-/-^and ID8-*Trp53*^-/-^; *Brca1*^-/-^cells. Log-rank (Mantel-Cox) test was applied to derive significant differences in (A). Mann-Whitney non-parametric test was used for (F-L) * p<0.05, ** p<0.005, *** p<0.001, **** p<0.0001. Gene expression data analysis was performed using nSolver Advanced Analysis Software. Mean ± SD.

Given our research question, all subsequent analyses were performed using the *Pten* deficient and *Brca1* deficient cell lines to investigate tumor immune associated disease states. I*n vivo* bioluminescence imaging revealed a more rapid tumor progression in mice implanted with *Pten* deficient cells compared to *Brca1* deficient cells by week 4 (Figure 2B). To investigate the effects of *Pten* or *Brca1* loss on the associated TIME, we characterized baseline immune transcriptome profiles of tumours generated from either *Pten* deficient or *Brca1* deficient cells (Figure 2C-2E; Supplemental Figure 1A-1B). A total of 119 genes were significantly differentially expressed between the two groups. NanoString nSolver based analysis revealed an enrichment in genes associated with innate and adaptive immunity, such as antigen processing and T cell function in tumors generated from *Brca1* deficient cells compared to those from *Pten* deficient cells (Figure 2C and 2D). Contrastingly, expression of genes associated with cancer progression, angiogenesis, and extracellular matrix stiffening (such as *Angpt2, Col3a1* and *Col1a1*) were significantly increased in *Pten* deficient tumours (Figure 2C and 2E). A 2-fold lower expression of IFN activated genes *Cxcl9* and *Cxcl10* was observed within *Pten* deficient tumors compared to the *Brca1* deficient tumors (Figure 2D). *Pten* deficient tumors also showed decreased expression of genes involved in several cytotoxic immune cells, however, genes associated with an exhausted T cell phenotype such as *Cd279* (PD-1) and *Lag3* were overexpressed in *Pten* deficient tumors (Supplemental Figure 2A).

Polychromatic flow cytometry-based analysis of both myeloid and lymphoid cells within the cellular fraction of ascites generated from *Pten* deficient cells showed significantly decreased proportions of CD8+ cytotoxic T cells (Supplemental Figure 1C) and activated CD8+CD69+ cytotoxic T cells compared to *Brca1* deficient ascites cells (Figure 2F). Significantly increased proportions of F4/80+ macrophages within the ascites generated from mice injected with *Pten* deficient cells was also observed (Figure 3B). A significant increase in suppressive immune populations, such as CD206+ M2 macrophages and GR1+ MDSCs was also observed in the ascites generated from *Pten* deficient cells (Figure 2G and 2H). Corresponding ascites cytokine levels showed elevated levels of macrophage chemoattractant protein 1 (MCP-1; Figure 3), IL-6 and IL-10 (Figure 2I; p = 0.017 and 2K; p = 0.029). Compared to ascites from mice injected with *Brca1* deficient cells, CXCL10, a chemokine critically involved in the recruitment of cytotoxic immune cells, level was significantly decreased in the ascites fluid from mice injected with *Pten* deficient cells (Figure 2L; p = 0.0314). Quantification of secreted chemokines from *Pten* deficient cells in supernatants derived from *in vitro* propagated cells revealed a similar trend with significantly decreased levels of both CXCL10 and CCL5 as compared to *Brca1* deficient cells (Supplemental Figure 1E and 1F; p=0.0022).

**Figure 3.**
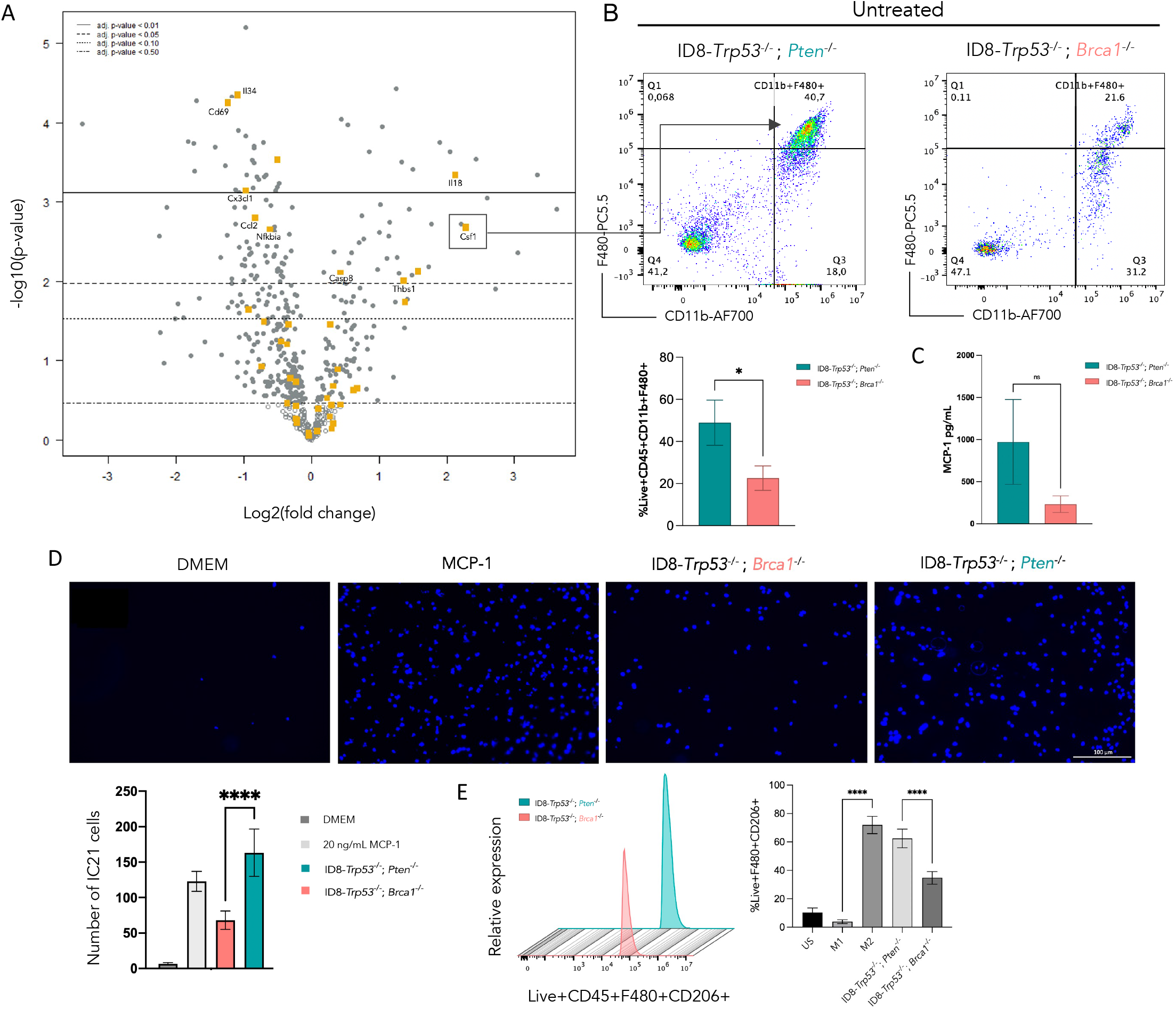
Pten deficient ovarian cancer cells alter their TIME towards a suppressive state via polarizing macrophages to a M2-like phenotype. Total RNA from untreated tumors of different genotypes was subject to NanoString gene expression profiling using the panCancer immune gene panel displayed as a (A) volcano plot showing differential expression pattern of genes associated with macrophage function. (B) Proportion of cells derived from ascites of mice injected with ID8-*Trp53*^-/-^; *Pten*^-/-^and ID8-*Trp53*^-/-^; *Brca1*^-/-^cells expressing CD45+CD11b+F4/80+ cells. (C) MCP-1 chemokine levels derived from the ascites fluid of untreated mice injected with ID8-*Trp53*^-/-^; *Pten*^-/-^and ID8-*Trp53*^-/-^; *Brca1*^-/-^cells. (D) Transwell migration assay of IC21 peritoneal macrophage cells stimulated (from left to right) with media control (DMEM), positive control (20 ng/mL MCP-1), or 72 h conditioned media of ID8-*Trp53*^-/-^; *Pten*^-/-^and ID8-*Trp53*^-/-^; *Brca1*^-/-^cells. Stimulated for 16 h; images 20x magnification. Flow cytometric analysis of IC21 macrophage proportions expressing (E) CD45+F4/80+CD206+ post-stimulation with concentrated ascites derived from mice injected with different genotypes of HGSC cells. US: unstimulated IC21 cells; M1 positive control represented by IC21 cells stimulated with IFN-γ (50 ng/mL) and LPS (100 ng/mL) in complete media; M2 positive control represented by IC21 cells stimulated with IL-4 (10 ng/mL) and IL-10 (20ng/mL) in complete media. Averages of triplicate wells/experiment for 3 repeated experiments displayed. Mann-Whitney non-parametric test was used for (B-C) * p<0.05 ** p<0.005, *** p<0.001, **** p<0.0001. One-way ANOVA applied for D and E. Mean ± SD. ns: not significant.

### *Pten* deficient ovarian cancer cells recruit and polarize macrophages into a M2-like phenotype

We next evaluated the expression profiles of genes associated with innate and adaptive immune cells in tumors from mice injected with *Pten* deficient or *Brca1* deficient ID8 cells. NanoString-based immune transcriptome analysis revealed increased expression of macrophage compared to *Pten* deficient tumors (Supplemental Figure 1G). While the expression of genes associated with macrophage phenotypes and function was higher in *Brca1* deficient tumors, significantly increased expression of genes such as colony-stimulating factor 1 (*Csf1;* p<0.05), responsible for the differentiation of hematopoietic stem cells into macrophages, was observed in *Pten* deficient tumors (Figure 3A).

While macrophage-associated intratumoral gene expression was relatively low in *Pten* deficient tumors compared to *Brca1* deficient tumors, flow cytometric analysis of ascites revealed significantly increased proportions of F4/80+ macrophages within the ascites generated from mice injected with *Pten* deficient cells (Figure 3B). Further, corresponding ascites cytokine levels showed elevated MCP-1 levels in *Pten* deficient mice (Figure 3C). Transwell migration assays of genetically distinct ID8 cells demonstrated a significantly increased migration of IC21 peritoneal macrophage cells incubated with conditioned media from *Pten* deficient cells compared to that from *Brca1* deficient cells (Figure 3D).

We next investigated whether cancer cell intrinsic loss of tumor suppressor genes influences the polarization of macrophages. Treatment of IC21 peritoneal macrophage cells with concentrated ascitic fluid derived from mice injected with *Pten* or *Brca1* deficient cells was used to assess polarization of IC21 macrophages into either an anti-tumor M1 or pro-tumor M2-like phenotype. A higher proportion of F4/80+CD206+ cells, indicating polarization to a M2-like phenotype, was observed upon treatment with ascites that was generated from *Pten* deficient mice (p=0.0079; Figure 3E). Of note, macrophages derived directly from the local ascites environment of mice injected with *Pten* deficient ID8 cells displayed a higher proportion of PD-L1 immune checkpoint expressing macrophages (Supplemental Figure 1D) compared to those from the *Brca1* deficient ascites. These results demonstrate that the suppressed tumor immune state associated with the loss of *Pten* in HGSC may be driven by increased polarization of tumor infiltrating macrophages into a M2-like phenotype.

### Host STING pathway is critical in anti-tumor immunity against *PTEN* deficient tumors

Given the established IFN-1 regulatory functions of *Brca1* and *Pten*, we next determined the cancer cell intrinsic activation of STING pathway. Immunofluorescence staining revealed increased cytosolic dsDNA in *Brca1* deficient cells compared to *Pten* deficient cells under unstimulated conditions (Figure 4A). Upon stimulation of *Pten* deficient or *Brca1* deficient cells with STING agonist, we also observed that STING protein levels in *Brca1* deficient cells peak at 3h post-treatment as compared to 6h for *Pten* deficient cells (Figure 4B). Moreover, expression levels of genes related to STING pathway, including *Tmem173 (STING; p>0*.*05), Irf3 (p=0*.*0355), NFκB (p=0*.*0013)*, chemokine *Cxcl10* (p>0.05) and *Ccl5 (p>0*.*05)*, and the receptors *Ifnar1, Ifnar2, and Ifngr1*, were decreased in *Pten* deficient tumors compared to *Brca1* deficient tumors (Figure 4C; Supplemental Figure 2A).

**Figure 4.**
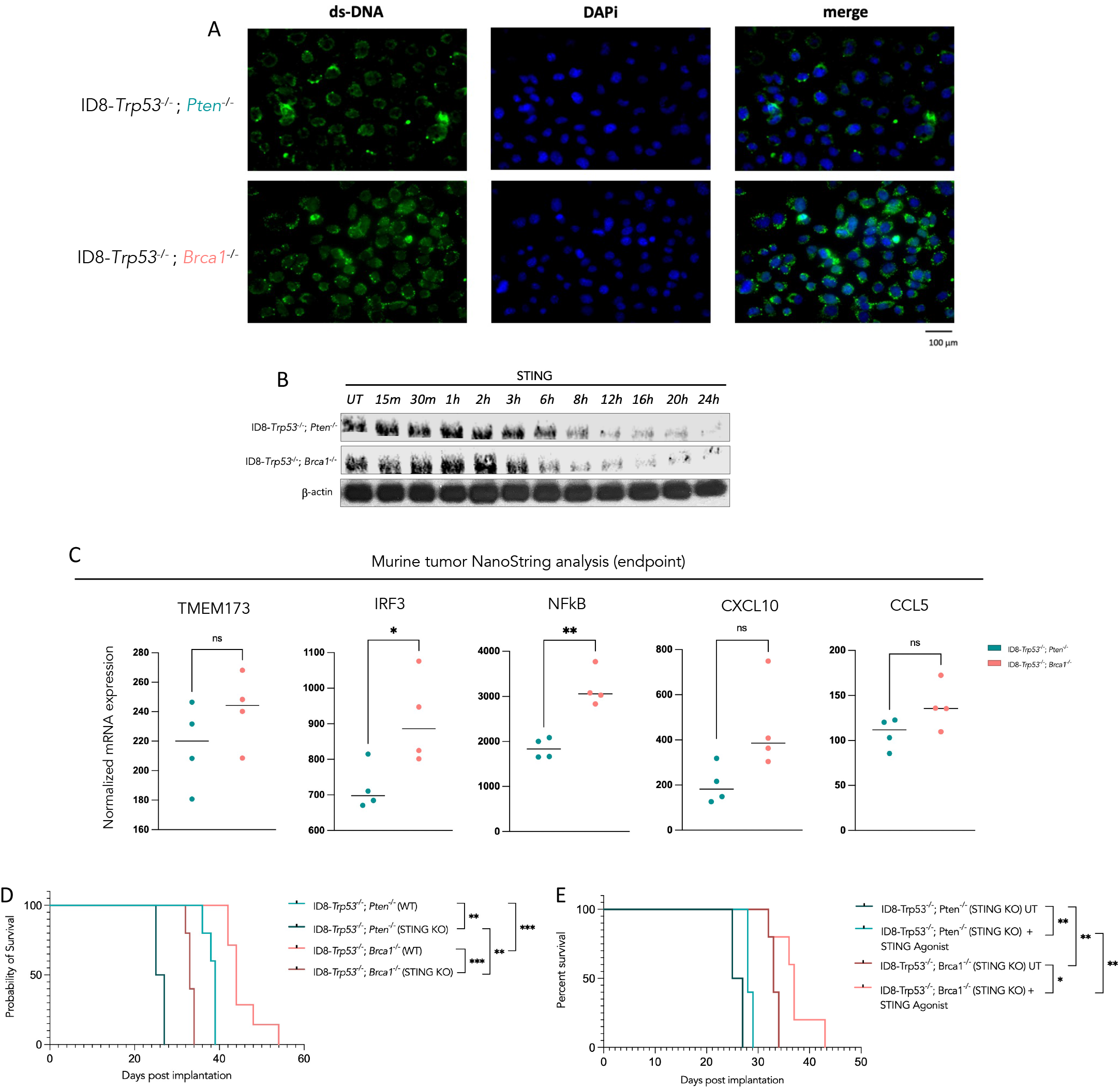
Host STING pathway activation is critical to improved response to chemotherapy. (A) Immunofluorescent confocal images for Immunofluorescent labelling of ds-DNA (FITC-green) and DAPi (nucleus, blue) in unstimulated ID8 cells of varying genotypes. Magnification, x40; scale bar 100 μm. (B) Peak activation of STING between cells of varying genotypes via assessing STING protein expression. (C) Isolated RNA from untreated tumors from ID8-*Trp53*^-/-^; *Pten*^-/-^and ID8-*Trp53*^-/-^; *Brca1*^-/-^cells were subjected to NanoString-based gene expression profiling to identify variation in genes associated with the cGAS-STING pathway. (D) Kaplan-Meier survival analysis of mice implanted with ID8-*Trp53*^-/-^; *Pten*^-/-^and ID8-*Trp53*^-/-^; *Brca1*^-/-^cells in wildtype (WT) C57BL6 mice or Sting1 deficient mice (STING-KO) was reported. (E) Kaplan Meier survival analysis of untreated control (UT) or STING agonist treated STING-KO mice injected with either ID8-*Trp53*^-/-^; *Pten*^-/-^and ID8-*Trp53*^-/-^; *Brca1*^-/-^cells. Log-rank test was performed to assess statistical significance between groups. (n = 5-15) for each treatment group. Mann-Whitney non-parametric test was applied for (C). * p<0.05 ** p<0.005, *** p<0.001, **** p<0.0001. Data analysis was performed using nSolver Advanced Analysis Software. ns: not significant.

We then investigated whether response to combination chemo-immunotherapy is affected by cancer cell intrinsic loss of *Pten* or *Brca1*. STING-KO mice injected with *Pten* deficient ID8 cells revealed significantly shorter median OS (26 days) compared to WT mice (median OS of 39 days; Figure 4D). Similar differences in survival were also observed in STING-KO mice injected with *Brca1* deficient ID8 cell compared to WT mice (median OS of 34 and 44 days, respectively; Figure 4D). This highlights the importance of host STING pathway in anti-tumor immunity independent of cancer cell intrinsic differences in STING pathway activation.

Further, the addition of exogenous activation of the STING pathway using a STING agonist resulted in no significant survival benefit in STING-KO mice injected with *Pten* deficient cells compared to the untreated mice (Figure 4E). However, treatment of STING-KO mice injected with *Brca1* deficient cells, with STING agonist significantly prolonged survival compared to the control group, suggestive of a role for pre-existing DDR deficiency-mediated increased IFN-1 response (Figure 4E).

### Exogenous activation of STING increases the response of Pten deficient tumors to carboplatin chemotherapy

Tumors exhibiting chemoresistant malignant cells, such as those with *Pten* deficiency, require improved response outcomes, which may be achieved by direct STING pathway activation. *In vitro* carboplatin chemosensitivity assay confirmed a higher IC50 in *Pten* deficient cells (3.755 μM) compared to *Brca1* deficient cells (0.6974 μM; Supplemental Figure 2B). Response to carboplatin chemotherapy was significantly increased in mice injected with *Pten* deficient ID8 cells following addition of STING agonist in the treatment regimen compared to those treated with carboplatin alone (median OS 64 and 51 days, respectively; Figure 5B). Kaplan-Meier survival analysis and log-rank (Mantel-Cox) test showed STING agonist monotherapy does not impart significant survival benefit in *Pten* deficient mice (median OS 38.5 days) compared to the vehicle group (median OS 39 days, *p*=0.0516; Figure 5B). Similar treatment response patterns were observed in mice injected with *Brca1* deficient cells (Figure 5C); however, the effects of prolonged survival were more pronounced in mice injected with *Pten* deficient ID8 cells (Figure 5B and 5C). Plasma CXCL10 and CCL5 cytokine levels were increased upon carboplatin and STING agonist treatment (Figure 5D and 5E), where levels of CXCL10 were higher in plasma of mice with untreated *Pten* deficient tumors compared to untreated *Brca1* deficient tumors.

**Figure 5.**
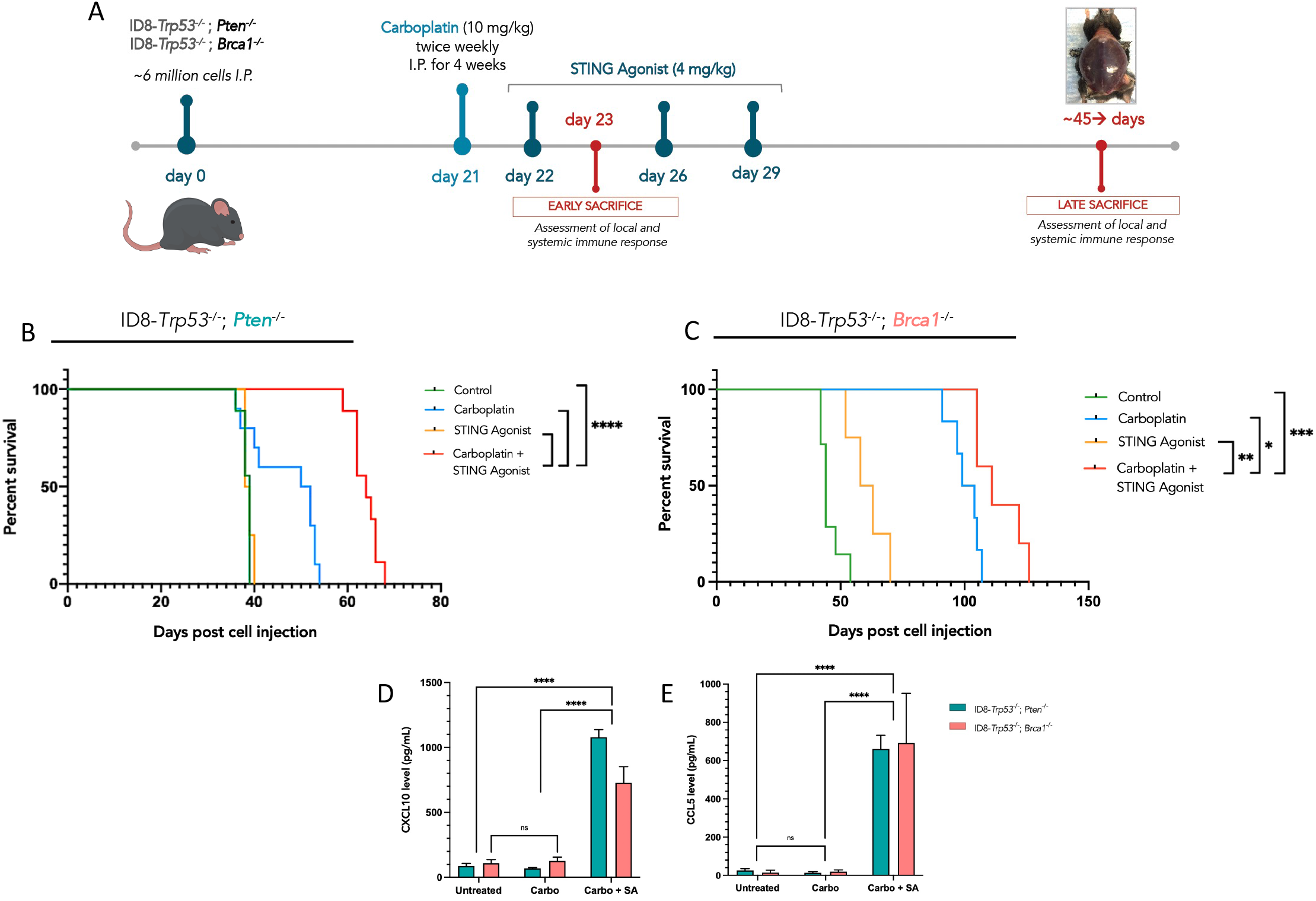
Treatment with STING agonist post carboplatin chemotherapy increases the response of Pten deficient ovarian tumors. (A) Schematic demonstrating the timeline of cell injection and subsequent treatment course and dosage of carboplatin and STING agonist. Kaplan-Meier survival analysis of mice injected with either (B) ID8*-Trp53*^-/-^; *Pten*^-/-^or (C) ID8-*Trp53*^-/-^; *Brca1*^-/-^cells for different treatment groups (n = 8-15 each). Cytokine profiles of (D) CXCL10 and (E) CCL5 from the plasma of untreated control (UT), carboplatin treated, or combination carboplatin and (+) STING agonist treated mice of either genotype. Log-rank (Mantel-Cox) test was applied to derive significant differences between treatment groups. * p<0.05 ** p<0.005, *** p<0.001, **** p<0.0001. Two-way ANOVA was applied. Mean ± SD.

### Exogenous STING pathway activation repolarized M2-like macrophages and increased T cell activation within the *Pten* deficient TIME

Based on the finding that STING pathway activation increased immune cell recruitment and imparted a survival benefit in *Pten* deficient tumors, we further investigated the mechanism by which this genotype-specific immune suppression occurs and the ability of STING activation to reinvigorate anti-tumor T cell activation. *In vitro* activated CD8+ T cells derived from age matched healthy mice revealed significantly decreased intracellular IFN-γ expression upon co-culture with macrophages from ascites of mice injected with *Pten* deficient ID8 cells compared to those from *Brca1* deficient ascites (p=0.008; Figure 6A). Indeed, *in vivo* treatment with STING agonist demonstrated a decrease in both myeloid proportions and a shift away from F4/80+ macrophages with a CD206+ M2-like phenotype, represented as a t-SNE plot of ascites from *Pten* deficient cells (Figure 6B). Additionally, proportions of CD69+CD8+ activated T cells in the ascites cellular fraction increased upon treatment with combination therapy compared to that of the vehicle treated mice (Figure 6C).

**Figure 6.**
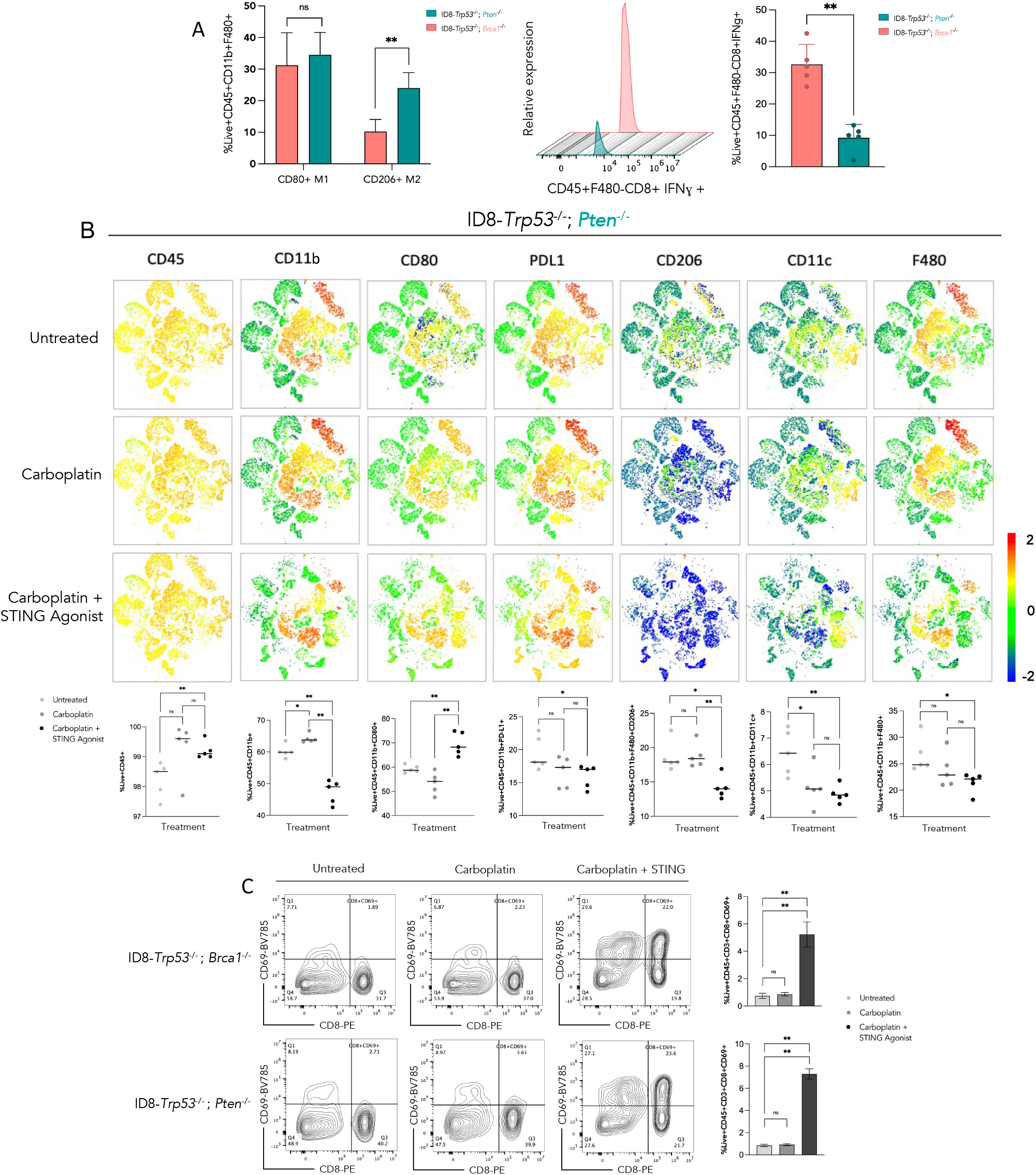
Exogenous STING pathway activation induces anti-tumor phenotype of immune cells within the Pten deficient TIME. (A) Co-culture of ascites-derived macrophages of mice injected with ID8*-Trp53*^-/-^; *Pten*^-/-^or ID8-*Trp53*^-/-^; *Brca1*^-/-^cells. Baseline proportions of CD80+ M1 macrophages and CD206+ M2 macrophages added to healthy T cells displayed. Histogram showing representative average of IFNγ producing CD8+ T cells from triplicate wells 48h post-stimulation with macrophages derived from ascites of either cell genotype (using n=5 mice for each genotype). Averages of triplicate wells used within this assay were displayed as a bar graph. (B) t-SNE plots generated from a concatenation of myeloid markers expressed on ascites derived from ID8*-Trp53*^-/-^; *Pten*^-/-^cells using Flowjo® FlowSOM and Phenograph Plug-in features (n=5 for each treatment group: untreated, carboplatin, and carboplatin + STING agonist). Corresponding cell proportions for each marker within ascites from individual mice displayed as bar graphs below. (C) Proportion of Live+CD45+CD3+CD8+CD69+ activated T cells within the ascites generated from untreated, carboplatin or carboplatin + STING agonist treated ID8*-Trp53*^-/-^; *Pten*^-/-^or ID8-*Trp53*^-/-^; *Brca1*^-/-^cells; analyzed using one-way ANOVA. Mann-Whitney non-parametric test was used for (A). * p<0.05 ** p<0.005, *** p<0.001, **** p<0.0001. Mean ± SD. ns: not significant.

## Discussion

Cancer cell intrinsic genetic alterations such as loss of tumor suppressor gene function via mutations, copy number alterations, or epigenetic modifications are key determinants of the pre-treatment TIME and clinical outcomes. The HGSC TIME exhibits a spectrum of immune active and underactive states that associate with differential treatment response. In the current study, we investigated this phenomenon in the BRCA1 and PTEN loss associated TIME to mimic the polarized HGSC disease states that exhibit contrasting immune landscapes. In both patient tumors and a syngeneic murine model of HGSC, we demonstrate how cancer cell intrinsic genetic alterations contribute to variable survival outcomes and tumor immune phenotypes. The *BRCA1* gene loss associated activation of STING pathway and consequent immune infiltration via induction of IFN-1 associated chemokines has been reported in ovarian cancer and other solid tumors^22^. However, the *PTEN* loss associated underactive TIME, chemoresistance and poor prognosis in HGSC remains understudied^23,24^. Findings from our study validated the previous reports^22,25^ on *BRCA1* deficiency associated cytosolic abundance of dsDNA in ID8 cells and downstream constitutive activation of the cGAS-STING pathway^22^. *PTEN*, however, functions to dephosphorylate IRF3 and stimulates its translocation into the nucleus to begin transcription of IFN-1 genes. Thus, its absence results in decreased cGAS-STING pathway activation and subsequent IFN-1 response.

While we previously demonstrated that chemoresistant HGSC patient tumors display a low density of CD8+ TILs and lower expression of IFN-1 genes^4^, the difficulty in treating HGSC lies primarily in our inability to precisely attribute the specific tumor immune states associated with vast genetic heterogeneity in this cancer. Indeed, widespread prevalence of *TP53* mutations in HGSC results in baseline DDR deficiency likely affecting IFN-1 response and innate immune activity via the cGAS-STING pathway^26^. Supporting this notion, recent *in vitro* studies in breast and pancreatic cell lines have demonstrated that *TP53* loss of function mutations, interfere with the cytoplasmic DNA sensing machinery required for cGAS-STING pathway activity^27^. Furthermore, mutations in *TP53* also correlate with increased expression of immune cell recruiting chemokines CXCL9, CXCL10 and CXCL11, and an inflammatory tumor immune microenvironment compared to wildtype *TP53* in HGSC specifically^28^. Here, we demonstrate that the effect of *Trp53* gene loss is further altered with concurrent mutations in DDR genes such as *Brca1* and *Pten*. Importantly, in this study we validated the relatively enhanced overall survival in mice injected with *Trp53* deficient ID8 cells alone, compared to those with additional loss of either *Pten* or *Brca1*, shedding light on the importance of determining the influence of paralleled loss of DDR genes on the TIME, tumor progression, and treatment responses.

In a cohort of adjuvant treated HGSC tumors, stratification of patients based on complete absence or presence of PTEN within both epithelial and stromal compartments revealed significantly decreased overall survival in patients with PTEN deficiency. Despite trends of decreased CD8+ T cells and CD68+ macrophages in the absence of PTEN, a high degree of variability was observed in this cohort, confirming the influence of cohort-specific differences on the challenges faced in biomarker translation within HGSC. Interestingly, PTEN deficient tumors displayed a significantly greater density of CD68+ macrophages within the stromal compartment compared to the epithelial compartment. Such differences were not observed in PTEN-intact tumors, suggesting a potential role for the loss of PTEN in the inhibition of immune migration into the tumor epithelial compartment. It is plausible that in tumors with PTEN loss, reduced infiltration of macrophages also impedes the process of tumor antigen cross presentation to CD8+ T cells. Their decreased chemokine expression in the TIME may thus further contribute to lower infiltration of immune cells contrasting tumors with *BRCA1* loss. Other mechanisms compounding this effect such as *PTEN*-loss associated hypoxia contribute to an “immune excluded” phenotype observed in other solid tumor cancers, may also be relevant within HGSC tumors^29^. A comprehensive understanding of intratumoral localization of macrophages and the associated environmental cues within the different tumor compartments on their resultant phenotype and function as M1 vs M2 is critical.

In mice injected with *Pten* deficient cells, both cellular and secreted profiles of malignant ascites and tumors revealed a suppressed and underactive tumor microenvironment. This was reflected by significantly decreased proportions of activated CD8+ T cells and increased GR1+ MDSCs. Similarly, genes associated with cancer progression, adhesion, and T cell exhaustion were overexpressed in *Pten* deficient ID8 tumors.

Interestingly, ascites from mice injected with *Pten* deficient cells showed significantly increased proportions of CD206+ M2 macrophages. *In vitro* analysis revealed *Pten* deficient cancer cell-secreted factors led to migration of significantly more IC21 peritoneal macrophages as compared to those from *Brca1* deficient cells. This suggests cancer intrinsic *PTEN* mutations promote an immune microenvironment with enhanced polarization of macrophages towards M2-like suppressive behavior. Such a polarization could also result from elevated levels of the cytokine IL-10, which was significantly higher in ascites from mice injected with *Pten* deficient cells. Another key finding reflecting an immunosuppressive microenvironment driven by *Pten* deficiency in tumors was the reduced expression of intracellular IFNγ in CD8+ T cells co-cultured with macrophages derived from ascites. Similar findings have been previously reported in colorectal cancer^30^. Thus, during *Pten*-deficient tumor progression and peritoneal metastasis, the pro-tumoral environment generated potentially impairs the effector function of critical cytotoxic immune populations relative to other genotypes.

Administration of intraperitoneal STING agonist was used primarily as a means to stimulate a potent IFN-1 response that would subsequently rescue or stimulate the function of pre-existing immune cells and increase the recruitment of additional anti-tumor immune cells via secretion of CXCR3 receptor binding chemokines. IFN-1s are well-established in their ability to impart robust anti-tumor immune effects via activation of antigen presenting cell priming of tumor antigens to T cells, subsequently leading to activation of cytotoxic T cells^31,32^. We previously established that exogenous STING pathway activation increased IFN-1 signaling by elevating levels of the immune cell recruiting chemokines CXCL10 and CCL5^18^, herein demonstrating increased CD8+ cytotoxic T cell recruitment in mice injected with either *Pten* deficient or *Brca1* deficient cells. Addition of STING agonist rescued the effector functions of immune cells within *Pten*-deficient HGSC tumors by increasing cytotoxic T cell activation, enhancing response to carboplatin, and resulted in a more pronounced increase in their overall survival. As such, HGSC tumors harbouring mutations resulting in underactive immune states or growth patterns, similar to that observed with *PTEN* deficiency, may be targeted for STING inducing therapies. The extensive genetic and immune heterogeneity in HGSC tumors makes it challenging to develop precise biomarkers of treatment response. However, a chemoresistant phenotype that exhibits a decreased pre-treatment IFN gene expression signature may benefit via incorporation of markers that combine genetic alterations and immune profiles, such as *PTEN* loss. A delayed activation of STING pathway in *Pten* deficient cells compared to *Brca1* deficient cells is also suggestive of additional IFN regulatory mechanisms leading to an ultimate effect of reduced chemokine secretion, which are beyond the scope of this study and require further investigation. In this study we also demonstrate benefit of cancer cell intrinsic STING activation demonstrated in *Pten* deficient cells, however, STING-KO (*Sting1*^-/-^) mice injected with *Pten* deficient cells resulted in decreased overall survival relative to WT C57BL/6 mice. The importance of host IFN-1 activation in cancer progression is evident, independent of genotype; further studies are required to gain an understanding on the role of STING pathway activation within immune cells, specifically myeloid cells in the context of *Pten* deficiency.

*PTEN* deficiency in tumors was associated with an abundance of immunosuppressive M2-like macrophages in the malignant ascites. Several mechanisms within the complex TIME of HGSC may indeed, contribute to an M2-dominated state. Most relevant to our study is the reduced feed-forward loop activation of STAT1 and CXCL10 that may act in parallel to other cancer cell intrinsic mechanisms^33,34^. In *BRCA1* deficient tumors, a significant active STAT1/CXCL10 axis in both the cancer and tumor infiltrating immune cells has been widely reported^22,35,36^. Furthermore, the decreased expression of CXCL10 by M2-like macrophages may be additive to the overall aggressive behavior of *Pten* deficient cancer cells^37^. Finally, M2-like TAMs are known to promote HGSC survival and angiogenesis^38^. Another key finding from our study is the increased proportions of PD-L1+ macrophages, which may inhibit the cytotoxic function of T cells. In mice treated with STING agonist, we observed a shift from M2 to M1 macrophages. This finding is in concordance with a similar recent study in both colorectal cancer and breast cancer models where repolarization of M2 macrophages to a M1 phenotype was observed upon exogenous STING pathway activation^30,39^. This suggests that STING pathway activation in *Pten* deficient HGSC displays the benefits associated with decreased M2s, such as decreased angiogenesis, secretion of immunosuppressive cytokines such as IL-10 and TGF-β, while providing rationale for the combined use of STING agonists and immune checkpoint blockade therapy, particularly PD-L1 antagonism. Although previous reports have revealed mechanistic links between *Pten* loss and macrophage driven cancers such as glioma, this remains to be investigated in HGSC and beyond the scope of current study. As such, tumors which exhibit inactive immune states resulting from genetic alterations such as *PTEN* loss may see greater benefit from direct STING pathway and IFN-1 activating therapies (i.e., via the use of STING agonists following an immunogenic cell death inducing chemotherapy, oncolytic viruses, etc.). Indeed, as recently reported, PARP inhibitors and other DDR kinase inhibitors have also been shown to indirectly activate the STING pathway albeit at a lesser magnitude.

Our study is not without limitations, particularly, our investigation focuses on the influence of individual and isolated genetic alterations. Human HGSC cells can express several co-occurring alterations^7^ and future work should incorporate these findings to further our understanding on the effects of the interactions and relationships of the several common gene losses reported in HGSC. Herein, we explore the effect of PTEN and BRCA1 loss in a metastatic syngeneic ID8 murine model of HGSC, however, investigating the impact of cancer cell intrinsic genotype on the resultant TIME within other models is critical to our understanding^40^. To conclude, activation of the STING pathway is largely influenced by pre-existing IFN regulatory genetic alterations such as DDR gene deficiency, prevalent in HGSC tumors. Exogenous activation of STING pathway overcomes the immunosuppressive local microenvironment driven by *Pten* loss in cancer cells. As further research considers the cancer cell intrinsic genetic alterations^41^, integrating genomic correlates of the TIME in HGSC patients will provide an improved platform for rapid evaluation of optimal therapeutic combinations.

## ACKNOWLEDGEMENTS

This work was supported by the Canadian Institutes of Health Research grant (CIHR, grant #159497), Ontario Ministry of Research Innovation and Science; Early Research Award and Queen’s University Research Initiation Grant to M. Koti. Additional support was provided to NS by the Franklin Bracken Fellowship, Dean’s Doctoral Award through Queen’s University, and the Ontario Graduate Scholarship program. We thank Shakeel Virk at the Queen’s Laboratory for Molecular Pathology (QLMP) for his assistance with TMA imaging and Halo, Katy Milne at BC Cancer’s Molecular and Cellular Immunology Core (MCIC) for immunofluorescence staining of TMAs and Natasha Jawa for support with R Studio.

## AUTHOR CONTRIBUTIONS

M. Koti designed the overall study, provided insights on experimental design and interpretation of results. NS designed and performed all experiments and analyzed the data. DL, and EL assisted with *in vivo* work. DL, GC, EL, JWS, AH, SC, and BL assisted with *in vitro* experiments. M. Köbel and KT contributed to human TMA based studies and analysis. KT provided guidance and assistance with overall statistical analysis. NS and M. Koti wrote the manuscript.

---

**Supplemental Table 1.** Description of clinicopathological features of patients with high-grade serous ovarian cancer.

| **Age (years)** |  |  |
| --- | --- | --- |
| Mean | 60.8 ± 10.08 |  |
| Median | 59 |  |
| Range | 44 – 92 |  |
| **Stage** | **Frequency** | **Percentage** |
| I | 1 | 0.91 |
| II | 11 | 10 |
| III | 91 | 82.73 |
| IV | 4 | 3.64 |
| Missing | 3 | 2.73 |
| **TOTAL** | 110 | 100 |

**Supplemental Figure 1.**
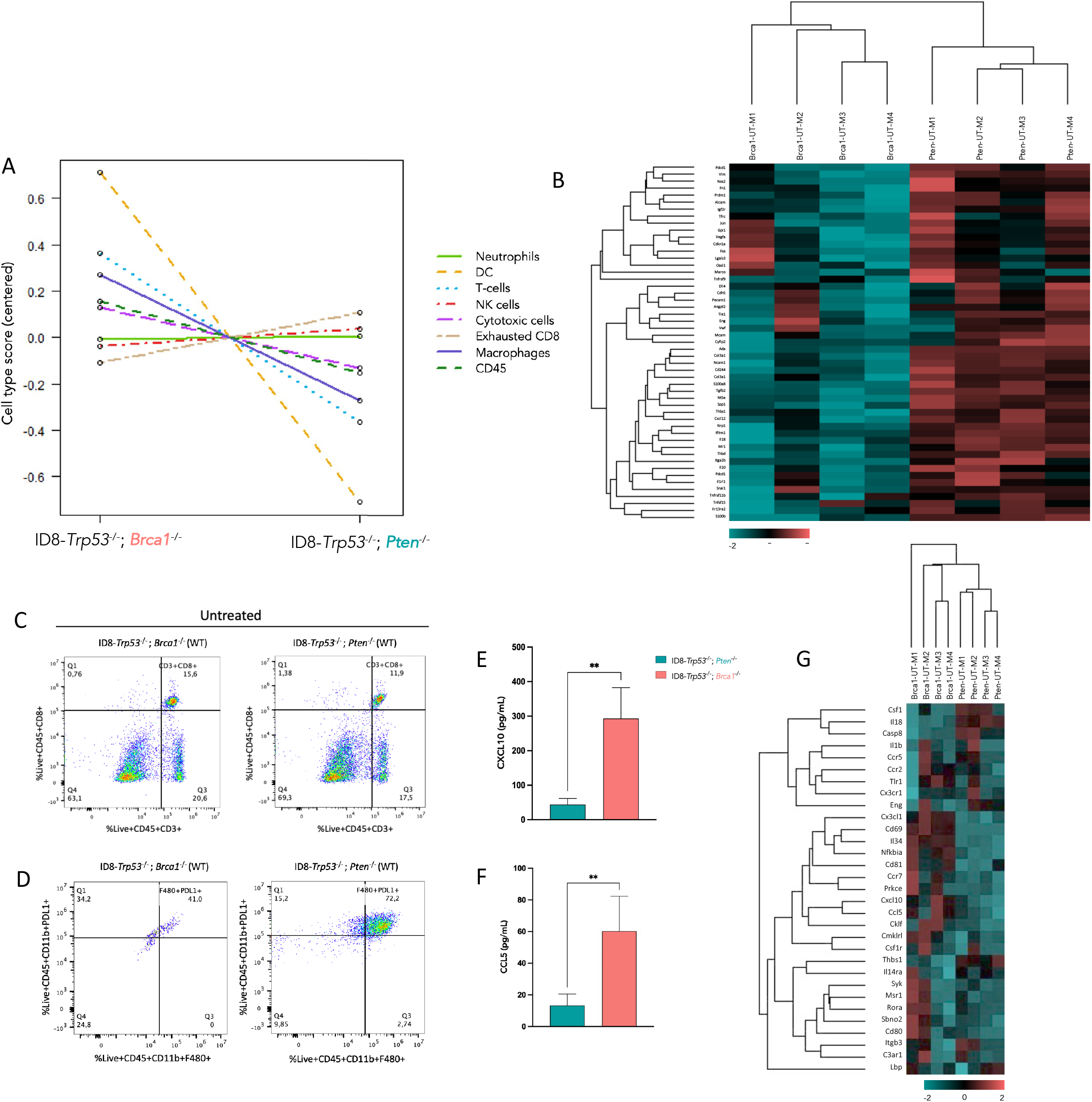
Pten deficient HGSC TIME and progression is distinct from Brca1 deficient HGSC. Total RNA from untreated tumors of different genotypes was subject to NanoString gene expression profiling using the panCancer immune gene panel displayed as (A) gene scores representing various immune cell types and (B) heatmap of 1.5-fold significantly differentially expressed genes between tumors generated from untreated ID8-*Trp53*^-/-^; *Pten*^-/-^and ID8-*Trp53*^-/-^; *Brca1*^-/-^cells using NanoString panCancer immune gene panel. Proportion of cells derived from ascites of mice injected with ID8-*Trp53*^-/-^; *Pten*^-/-^and ID8-*Trp53*^-/-^; *Brca1*^-/-^cells expressing (C) CD45+CD3+CD8+, (D) CD45+CD11b+F480+PDL1+. (E) CXCL10 and (F) CCL5 cytokine levels within 48-hour conditioned media from ID8-*Trp53*^-/-^; *Pten*^-/-^and ID8-*Trp53*^-/-^; *Brca1*^-/-^cells. (G) Heatmap displaying macrophage associated gene expression between ID8-*Trp53*^-/-^; *Pten*^-/-^and ID8-*Trp53*^-/-^; *Brca1*^-/-^derived tumors. Cytokine experiments were derived from duplicate experiments of triplicate wells. Data analysis was performed using nSolver Advanced Analysis Software. Mann-Whitney non-parametric test was used. * p<0.05 ** p<0.005, *** p<0.001, **** p<0.0001. Mean ± SD. ns: not significant.

**Supplemental Figure 2.**
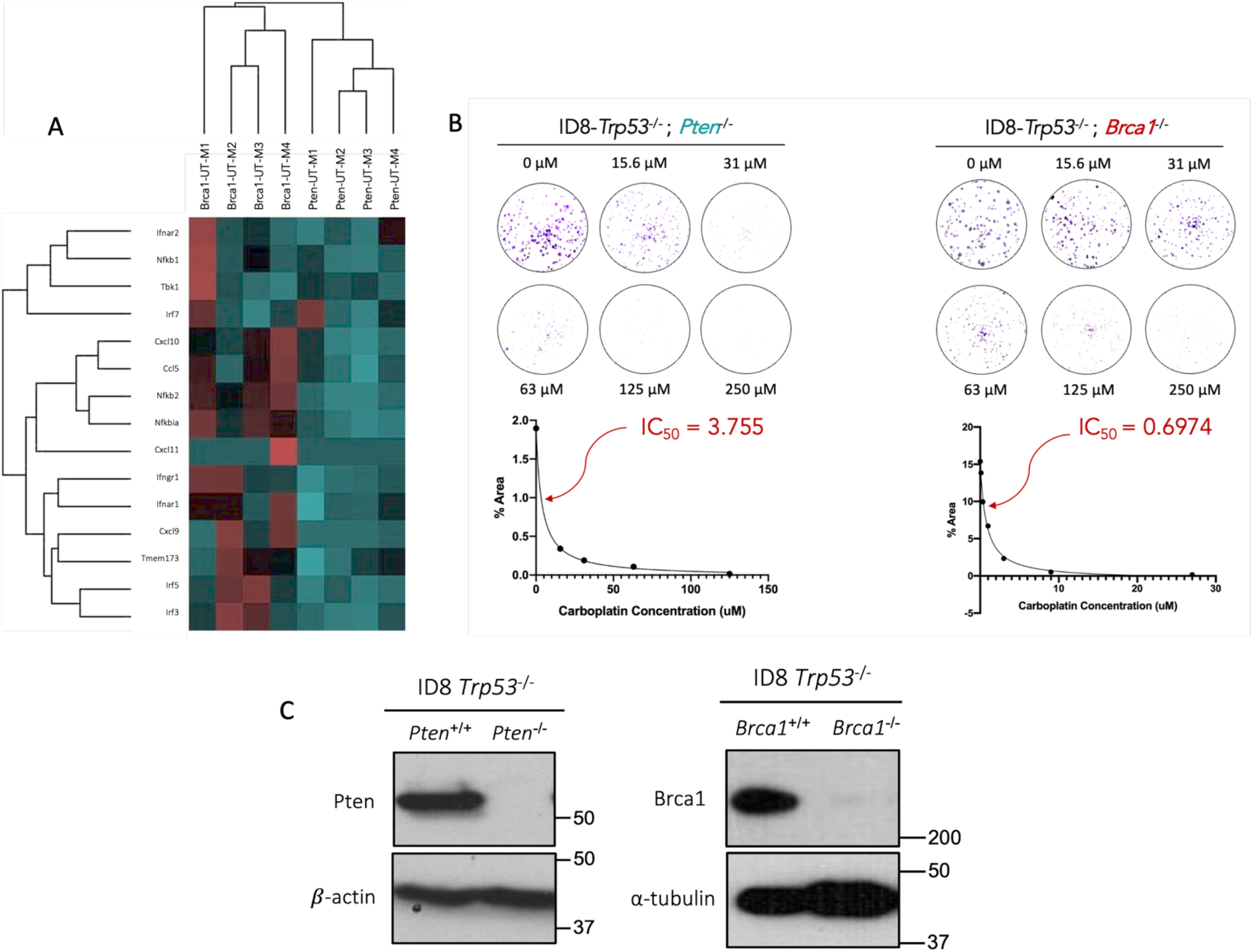
Host STING pathway activation is critical to improved response to chemotherapy. Total RNA from untreated tumors generated from genetically distinct ID8 cells displayed as a (A) heatmap displaying 1.5-fold differentially expressed STING pathway associated genes between tumors generated from untreated ID8-*Trp53*^-/-^; *Pten*^-/-^and ID8-*Trp53*^-/-^; *Brca1*^-/-^cells. (B) IC50 of carboplatin for ID8-*Trp53*^-/-^; *Pten*^-/-^and ID8-*Trp53*^-/-^; *Brca1*^-/-^cells. (C) Immunoblotting to assess for complete KO of Pten and Brca1 in cell lines prior to use. Data analysis was performed using nSolver Advanced Analysis Software.

**Supplemental Figure 3.**
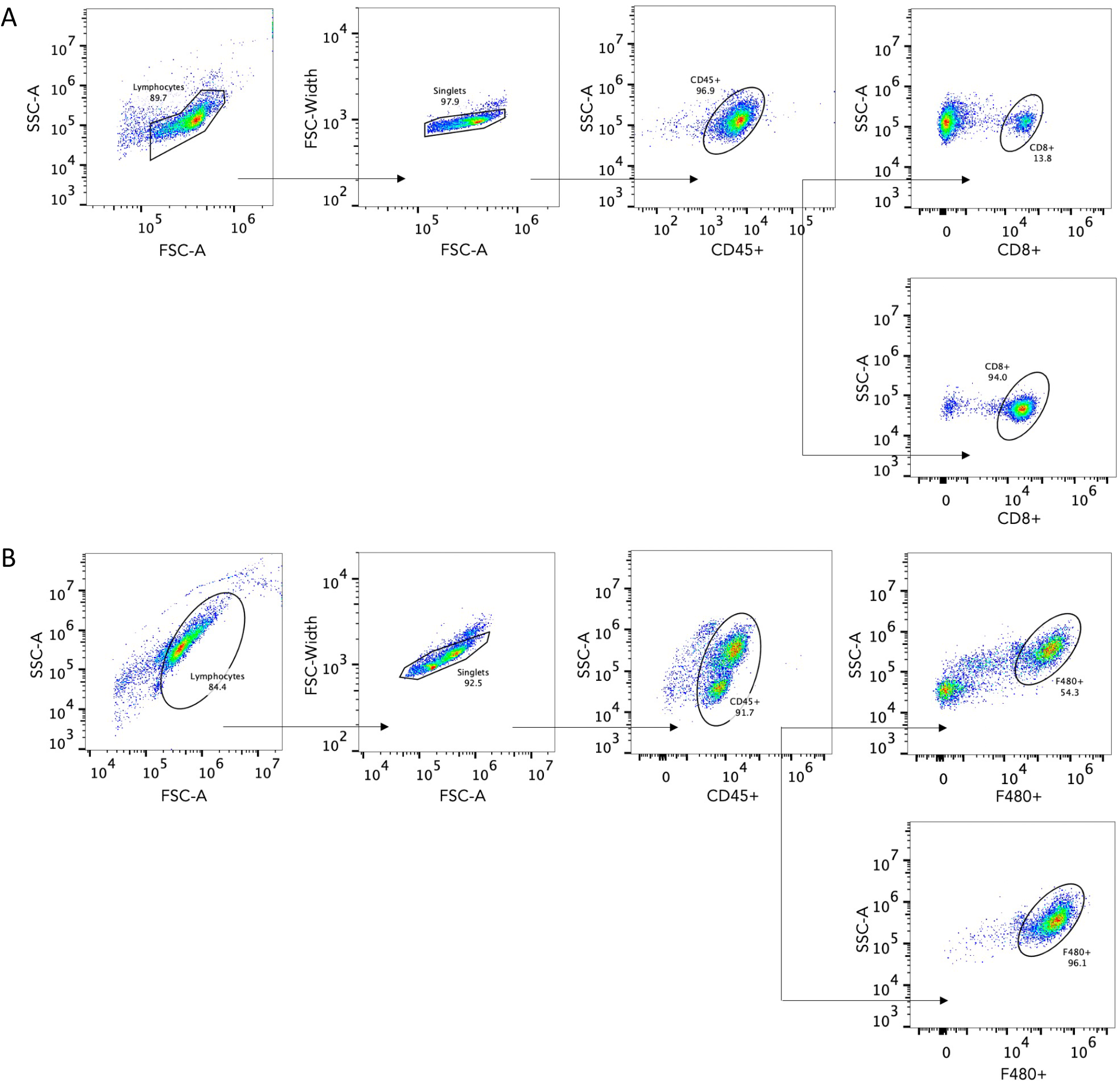
Isolation of immune cell proportions of CD8+ cytotoxic T cells from healthy spleen and macrophages from ascites. Flow cytometric analysis of immune cell proportions revealing proportion of isolated (A) CD8+ cytotoxic T cells and (B) F480+ macrophages used for T cell-macrophage co-culture assay assessing IFNγ producing T cells after exposure to macrophages derived from ascites of ID8*-Trp53*^-/-^; *Pten*^-/-^or ID8-*Trp53*^-/-^; *Brca1*^-/-^injected mice.

